# Asymmetric evolvability leads to specialization without trade-offs

**DOI:** 10.1101/2020.09.11.264481

**Authors:** Jeremy A. Draghi

## Abstract

Many ideas about the evolution of specialization rely on trade-offs—an inability for one organism to express maximal performance in two or more environments. However, optimal foraging theory suggests that populations can evolve specialization on a superior resource without explicit trade-offs. Classical results in population genetics show that the process of adaptation can be biased toward further improvement in already productive environments, potentially widening the gap between superior and inferior resources. Here I synthesize these approaches with new insights on evolvability at low recombination rates, showing that emergent asymmetries in evolvability can push a population toward specialization in the absence of trade-offs. Simulations are used to demonstrate how adaptation to a more common environment interferes with adaptation to a less common but otherwise equal alternative environment. Shaped by recombination rates and other population-genetic parameters, this process results in either the retention of a generalist niche without trade-offs or entrapment at the local optimum of specialization on the common environment. These modeling results predict that transient differences in evolvability across traits during an episode of adaptation could have long-term consequences for a population’s niche.

## Introduction

Species have limits that emerge from complex evolutionary processes. The breadth of a niche is proscribed by competition with other species, but also by constraints internal to the population: for example, limits to local adaptation in the face of gene flow (Kirkpatrick & Barton 1997). Variation in niche breadth is a key dimension of biodiversity, and is typically quite high (Poisot et al. 2015) but is declining as specialists are disproportionally at risk due to environmental change (Clavel et al. 2011). Understanding how organisms evolve to be specialists and what causes a niche to expand or contract is major goal of evolutionary ecology. While these questions are long-standing, evolution experiments focusing on niche breadth or host range, often in microbes (Kassen 2002), have helped to highlight the many gaps in how we think about evolution of the niche (Bono et al. 2020).

One highly influential explanation for restricted niche breaths is that they are limited by trade-offs, such that specialists can exceed the performance of generalists where their niches overlap. Despite the intuitive appeal of trade-offs, empirical evidence that they actually explain niche breadths has been hard to find (Futuyma & Moreno 1988; Fry 1996; Remold et al. 2012). This gap has motivated the search for alternative theories that incorporate behavior, population genetics, and macroevolution into a synthetic theory of niche limits (Poisot et al. 2011; Sexton et al. 2017).

A major insight from population-genetic models of niche breadth is that the evolution of preference for a particular habitat, and performance in that habitat, can be linked in a positive feedback loop (Holt 1985; Crespi 2004). Such models elaborate upon optimal foraging perspectives by allowing the value of resources to change with an organism’s degree of local adaptation, with these changes feeding back to determine which preferences are optimal (Futuyma & Moreno 1988). The cause of this feedback is that the intensity of selection to maintain and improve performance in a specific habitat is proportional to the number and reproductive success of organisms within it (Holt & Gaines 1992). This concept is identical to the more familiar example of weakening selection with increasing age due to declining reproductive value of later age classes; essentially, rarely encountered or unproductive environments are similar to rarely attained or less fecund age classes (Holt 1996a). If organisms evolve to prefer a habitat that confers greater reproductive success, then the accompanying shift in the focus of selection may, over time, further boost their fitness in the favored habitat. This prediction that stronger selection improves the degree of adaptation is quite general, emerging from models of trade-offs, but also from models where performance in each environment depend on independent sets of traits (e.g., Kawecki et al. 1997). Improved adaptation in one environment can then drive still greater levels of preference, completing the feedback loop. In such models, specialization can therefore arise from even minor asymmetries in performance across a broad niche (Fry 1996).

Any model of niche breadth that considers both performance traits and habitat selection incorporates this positive feedback loop. Such models are plentiful and highly diverse (see reviews in Futuyma & Moreno 1988; Wilson & Yoshimura 1994; Ravigné et al. 2009), but most inherit the idea from Levins (1962) of representing trade-offs with, in two dimensions, a curve of the maximum fitness in one environment given each possible fitness in the other environment (essentially, a Pareto front—see Shovel et al. 2012). Evolution of the degree of specialization can then be envisioned as movement in a single dimension, constrained to this curve of optimal combinations of fitnesses. Modeling two dimensions of performance in a single variable is an appealing simplification, but cannot adequately represent a scenario in which a population is suboptimally adapted to both environments. This scenario might arise when an invasive species adapts to the multiple novelties of a new environment (Prentis et al. 2008) or as a result of antagonistic coevolution with multiple hosts or prey species (e.g., Hall et al. 2011). In these circumstances, the rate of adaptive evolution, or evolvability, in each environment can now dynamically shape the value of each environment for reproductive success, and could therefore drive evolution of preferences for those environments.

Motivated by mixed evidence for trade-offs, a number of papers have explored the idea that generalists may suffer deficits in evolvability, even when trade-offs are avoided by virtue of separate genetic bases for performance in each environment (Kawecki 1994; Fry 1996; Holt 1996b; Whitlock 1996). Because generalists experience selection in multiple environments, the strength of selection for performance in any one of those environments is weaker in comparison to a specialist population that only reproduces in that single environment. This is fundamentally the same idea as the positive feedback between preference and performance discussed above, and this connection has led to predictions that, without trade-offs, specialists may replace generalists because they can adapt faster (Kawecki 1998) or because generalists cannot maintain fitness for rare environments in the face of deleterious mutation (Kawecki et al. 1997). While intriguing, these findings preceded the development of our modern concept of evolvability and its determinants, and haven’t shown that specialists can evolve with both the absence of trade-offs and the presence of substantial ecological costs of specialism, such as the search costs which are typically associated with a narrow niche in classical models of optimal foraging (Charnov 1976; Rosenzweig 1981).

Here, I show that specialization can evolve as initially generalist populations adapt to improve fitness in two environments, and that this adaptive specialization can occur without trade-offs and with substantial search costs imposed on specialists. Populations can evolve a reduction in their niche that is adaptive, in the sense that a sequence of beneficial mutations can lead a population to a local optimum of specialism. However, the fitness landscape allows for no-cost generalists that are competitively superior to any specialist; specialism therefore represents a local, inferior optimum. A population’s chance of adapting to this local optimum, and therefore specializing on the more common environment, is primarily driven by the relative evolvability of its performance in each environment, with recombination between the genetic bases of each trait emerging as a major determinant of whether populations retain generalism.

Recombination helps maintain generalism because it alleviates a type of clonal interference—competition between beneficial mutations that are simultaneously polymorphic (Gerrish & Lenski 1996)—that further impedes adaptation in traits with already low evolvability. Recent theory focusing on asexual evolution has predicted that adaptation among sites with small selection coefficients can be effectively stalled by interference from rapidly evolving sites with larger effects (Schiffels et al. 2011). Gomez et al. (2020) show that a trait with more frequent or larger beneficial mutations can effectively stall adaptation in a trait with lower evolvability, an effect that Venkataram et al. (2019) recently demonstrated experimentally. Here I apply these ideas to differences in evolvability that arise from variation in the frequency of two environments, with no other intrinsic differences in their quality. This scenario represents a challenging case for specialization, as both resources have identical initial and potential values. Specialization is seen to evolve when the less common environment becomes unprofitable in comparison to the more common environment, solely because of slow relative improvement in how organisms can exploit the rare environment. These results demonstrate the value of synthesizing our emerging understanding of evolvability with classical eco-evolutionary questions about the evolution of the niche.

## Model

### Overview of Ecology & the Life Cycle

The basic Wright-Fisher model was modified to allow for a potentially costly preference for the common environment, while retaining several traditional components of the model: non-overlapping generations and construction of a fixed number of adults via random sampling from an unlimited pool of gametes, producing approximately Poisson-distributed variation in reproductive success. Each organism has three traits—a preference trait (*f*) and two performance traits, w_A_ and w_B_—that are entirely determined genetically by three sets of distinct loci. The number of adults is limited to a carrying capacity *K*; each of these individuals then attempts to settle in a habitat of either environment A or B, as described below. The expected fecundity of an individual *i* in environment *x* is then:

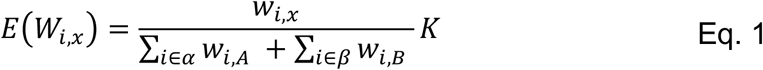

As implied by Eq. 1, the preference trait has no direct effect on fitness. Here *α* is the set of individuals that have settled in habitats of environment A, and *β* is the set in environment B. Because of search costs, as detailed below, |α| + |β| ≤ *K*; that is, the total number of individuals settled in each environment is no greater than *K*, but can be less than that. Following the taxonomy summarized in Ravigne et al. 2009, this is a model with global regulation of population size and variable habitat outputs; it is therefore a model of ‘hard selection,’ which generally is more likely to permit the success of specialists (Futuyma & Moreno 1988).

In fully asexual simulations, the fecundities given by Eq. 1 are then used as weights in a multinomial distribution from which *K* parents are selected, with replacement, to reproduce and form the next generation. In simulations with recombination, 2K parents are drawn with replacement, and offspring are determined via the linkage relationships described below. To avoid confounds in comparing asexual and sexual populations, all organisms modeled here are haploid.

### Environment Preference & Specialism

For simplicity, I focus here on scenarios in which environment A is more commonly encountered than environment B, and in which the preference trait causes aversion to the rarer environment (and therefore, a preference for the common environment A). The preference trait *f* is therefore treated as a probability to reject environment B when encountered, with a value of zero representing unbiased generalism and a value of one representing complete specialization on environment A. While it would be straightforward for future work to extend this model to allow preferences for either environment to evolve, this complication isn’t necessary to model the process of specialization on the common environment. Therefore, the model is designed asymmetrically and does not allow an environment-B specialist to evolve.

Each adult searches for a habitat, encountering environment A with probability 1 - *p* and environment B with probability *p*, where *p* < 0.5. These encounter probabilities are independent of genotype. If *f* > 0, then an individual will reject environment B with probability *f* and then experience a cost of searching, *c*, representing the probability of death while searching for a new habitat. If the individual avoids death it once again may encounter environment A with probability 1 - *p* or environment B with probability *p*, and may again reject B with probability *f*; this search process continues until each individual has perished or been assigned to an environment. The probabilities of assignment to A, B, or death are given by the sums of geometric series as follows:

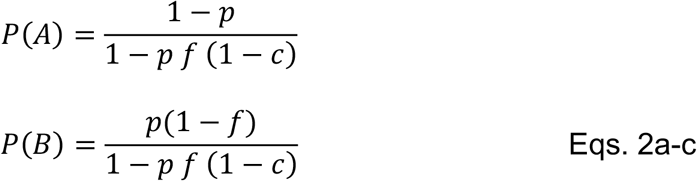

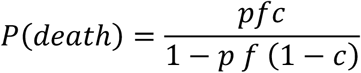

This model of search costs is similar to Forbes et al. (2017), but is generalized to allow a probabilistic preference.

Mean fitness of a given genotype can then be written as a sum of Eqs. 2a and 2b, weighted by that genotype’s fitness in those environments.

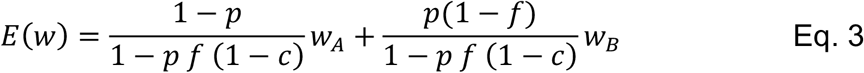

The derivative of Eq. 3 with respect to *f* at *f* = 0 is positive when the following inequality is met:

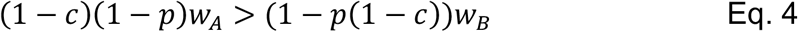

We can therefore predict that specialization begins to be favorable when Eq. 4 is true.

Two simplifications of Eq. 4 help establish some intuition. When *c* is zero, then specialism on A can start to evolve whenever *w*_*A*_ > *w*_*B*_. When, instead, *p* is small we can approximate the condition in Eq. 4 as (1 − *c*)*w*_*A*_ > *w*_*B*_; that is, specialism starts to be favorable when it is more profitable to risk death than to settle for environment B.

### Genetic Bases of Traits & Mutation

Fitness in environment A, *w*_A_, is determined by a set of loci *G*_A_. This set consists of *L* linked loci, each with two possible alleles and each with a fitness difference of *s*_A_ between them. Fitness effects combine multiplicatively across loci. Fitness in environment B is determined in exactly the same way, by an independent set of *L* loci *G*_B_ with fitness difference *s*_B_ between the alleles at each locus. *G*_A_ has no effect on fitness in environment B and vice versa. Within a set each locus is interchangeable with the others, so we can fully represent the genotypic basis of fitness for, say, environment A, by the number of beneficial alleles, *g*_A_, it contains.

A preference locus, represented by a single number in [0,1], is also encoded in each genotype. In simulations with linkage, all these elements are considered to be on the same chromosome in the order of the preference locus, *G*_A_, and *G*_B_. This is clearly not realistic for polygenic traits; the model is best seen as representing complete linkage with specific, purposeful deviations from that pattern. Linkage between *G*_A_ and *G*_B_ is a focus of the model, while linkage between the preference locus and *G*_A_/*G*_B_ is a nuisance factor—therefore, in some simulations the preference locus is modeled as if it is on an independent chromosome to assess the impact of this choice.

Mutation of the preference locus is implemented by simply redrawing the value from a uniform distribution. For *G*_A_ and *G*_B_, an additional parameter, *b*, is introduced that reflects a bias in favor of deleterious mutation. Smaller values of *b* decrease the chance that mutation will produce the beneficial allele. The probability that a mutation in *G*_A_ is beneficial (increasing *g*_A_) is 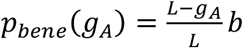, the probability that it is deleterious (decreasing *g*_A_) is 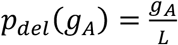, and remaining mutations are neutral. As a consequence, the rate at which new beneficial mutations arise decreases linearly with the degree of adaptation.

The number of mutations in each trait in a generation is drawn from a Poisson distribution with a mean of μ_pref_*N* for the preference trait, μ_A_*N* for *G*_A_ and μ_B_*N* for *G*_B_. Individuals are selected to receive mutations with replacement.

### Density dependence

In the absence of any habitat preferences, we expect (1-*p*)*K* adults to reproduce in environment A and *pK* in environment B. We refer to these values as the neutral expectations for densities in each environment. I implemented density-dependence growth as a discrete option: either density in each habitat was ignored, as described above, or a density higher than the neutral expectation for that environment caused reduced reproduction for everyone in that environment. The reduction in growth was calculated to be proportional to the percentage by which density exceeded the neutral expectation. If N_A_ is the number of breeding adults in environment A and adults encounter environment A with probability *p*, then reproduction in environment A was divided by a factor N_A_/(1-p)K whenever N_A_ > (1-p)K. If N_A_ did not exceed (1-p)K then fitness was calculated normally—there was no positive effect of low density. The corresponding calculation was also performed for environment B; by definition, only one environments could suffer negative density dependence in a given generation.

### Simulation approach

Aside from Eq. 4, all other results are the products of individual-based simulations. Simulations were written in R with integrated C++ algorithms for the sake of speed. In general, between 250 and 1000 replicates were performed for each treatment; confidence intervals are included in all figures in which they are not negligible. All code will be made publicly available upon publication.

**Table 1:**
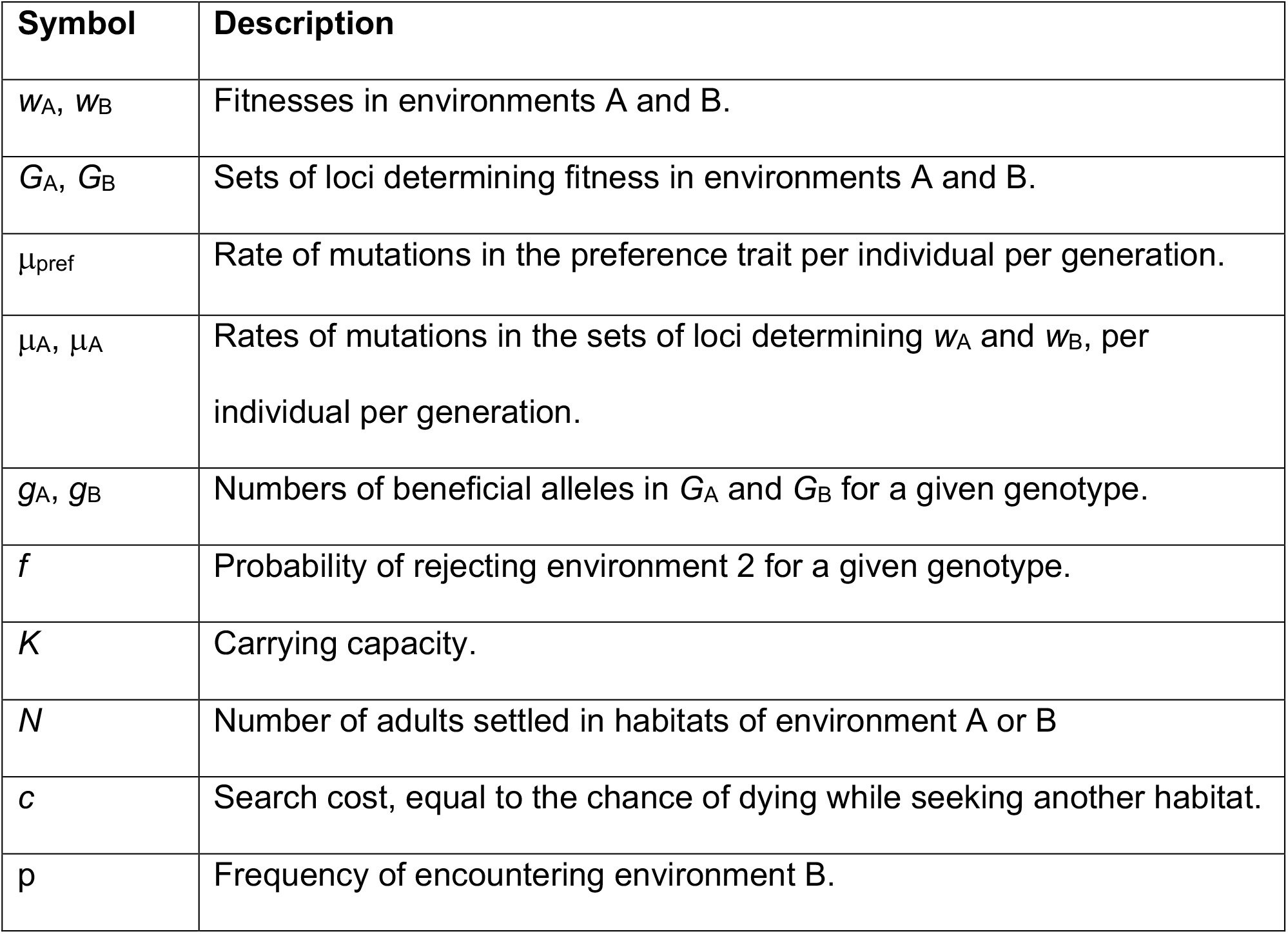
Important parameters.

| Symbol | Description |
| --- | --- |
| $w_A, w_B$ | Fitnesses in environments A and B. |
| $G_A, G_B$ | Sets of loci determining fitness in environments A and B. |
| $\mu_{\text{pref}}$ | Rate of mutations in the preference trait per individual per generation. |
| $\mu_A, \mu_B$ | Rates of mutations in the sets of loci determining $w_A$ and $w_B$, per individual per generation. |
| $g_A, g_B$ | Numbers of beneficial alleles in $G_A$ and $G_B$ for a given genotype. |
| $f$ | Probability of rejecting environment 2 for a given genotype. |
| $K$ | Carrying capacity. |
| $N$ | Number of adults settled in habitats of environment A or B |
| $c$ | Search cost, equal to the chance of dying while seeking another habitat. |
| $p$ | Frequency of encountering environment B. |

## Results

### Linkage between the genetic bases of traits promotes adaptive aversion to the rare environment

To understand how performance in each environment was shaped by adaptation to the other, I first performed simulations in generalists that could not evolve into specialists (i.e., μ_pref_ = 0). In large populations (*K* = 10,000) in which environment B was encountered by 25% of individuals (*p* = 0.25), the initial founders had with equal fitnesses in each environment. Performance in each environment was determined by a distinct set of one hundred loci (*L* = 100) for each environment. At each of these two hundred loci, one of two alleles conferred a small, multiplicative benefit (*s* = 0.01) in the corresponding environment. For the set of loci corresponding to environment A, all alleles are neutral in environment B, and vice versa. Therefore, there are no genetic trade-offs in performance across the environments, and no-cost generalists are possible. The starting genotype had an initial fitness of approximately 0.5 in each environment. Populations were then allowed to improve performance in one or both environments, by setting the mutation rates to (μ_A_ = 5 × 10^−4^, μ_B_ = 0), (μ_A_ = 0, μ_B_ = 5 × 10^−4^), or (μ_A_ = 5 × 10^−4^, μ_B_ = 5 × 10^−4^).

Figure 1 plots mean fitness in each environment across four hundred replicates, comparing fitness in each environment when that trait evolves alone (no genetic variation in the other trait) versus when it evolves simultaneously with the other fitness trait. Adaptation to the more common environment (environment A) is unaffected by simultaneous adaptation to the less common environment. However, with complete linkage (r = 0), adaptation to the less common environment is greatly slowed when the population is also adapting to environment A; this slowdown is largely but not entirely eliminated with free recombination between the genetic bases of each trait, *G*_A_ and *G*_B_. (Loci within each block are still completely linked). Recombination does not eliminate all interference between traits because the strength of selection in an environment is still dependent on that environment’s contribution to the total pool of offspring; *w*_A_ increases faster than *w*_B_ because p < 0.5, leading to weakening selection on *w*_B_ as the simulation progresses.

**Figure 1:**
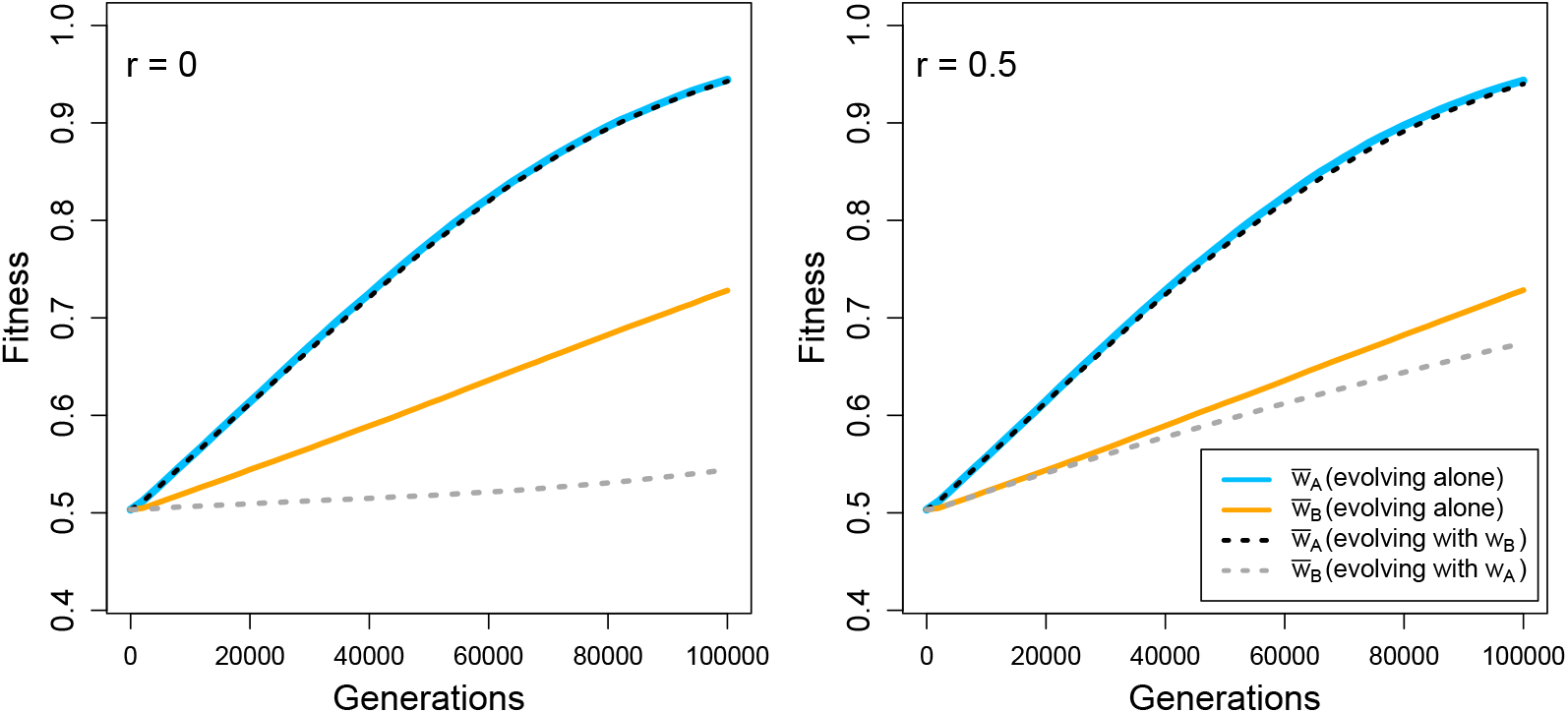
Mean fitness in each environment when each performance trait evolves alone (solid lines) or together (dotted lines). *Left*: No recombination between the sets of loci for performance in each environment (*r* = 0). *Right*: Full recombination between sets of loci (but still complete linkage within each set of loci). Four hundred replicates are averaged; 95% confidence intervals (not plotted) span less than 0.01 units on the *y*-axis. *K* = 10,000, *L* = 100, mutation per locus is either 5 × 10^−4^ (when allowed to evolve) or zero (when not allowed to evolve).

To understand how these asymmetries in evolvability interact with the evolution of preference, I performed simulations in which the preference trait was also allowed to evolve. Figure 2 shows two depictions of the same four representative examples of simultaneous evolution of all three traits. Fig. 2A plots the ratio of mean fitnesses, 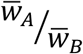, as both adapt. Evolution of the preference, depicted with the changing colors in Fig. 2, is predicted by the horizontal dashed line drawn from Eq. 4. Fig. 2B shows the same data with each performance trait shown on its own axis, with the line defined by Eq. 4 now appearing as a diagonal line.

**Figure 2:**
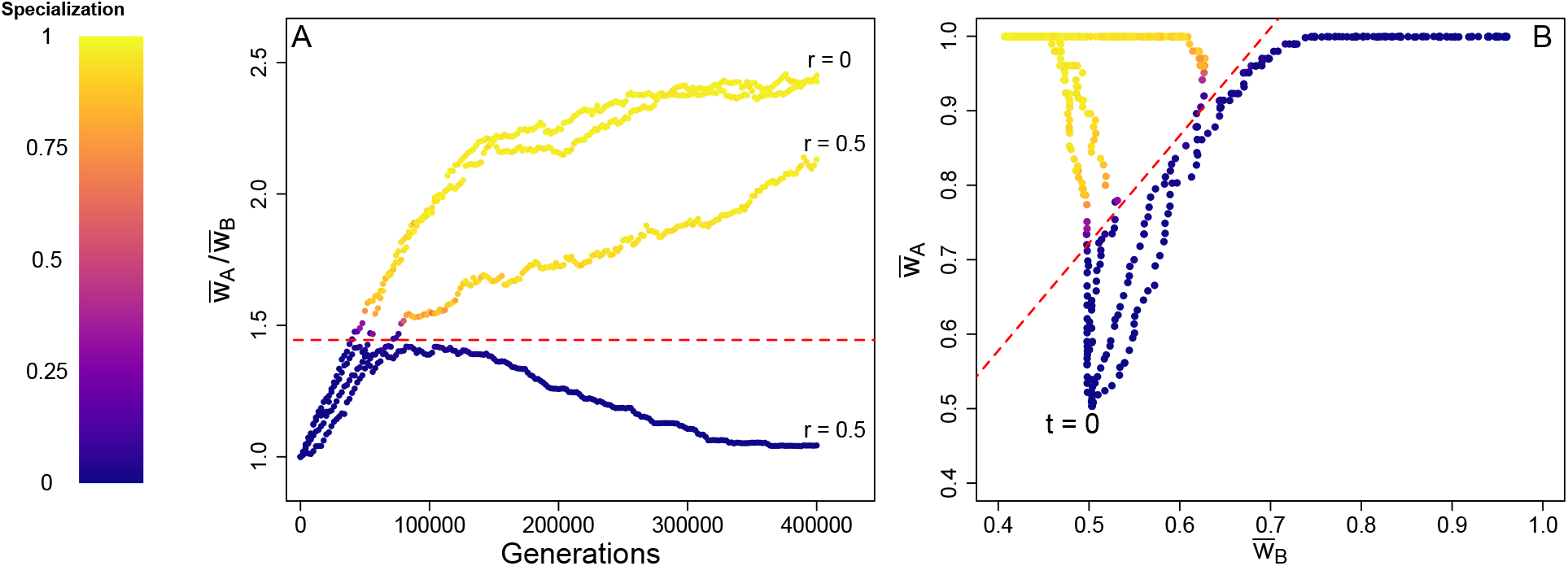
Representative samples of fitness evolution with complete linkage (*r* = 0) and free recombination between the genetic bases of fitness in environment A and B (*r* = 0.5). Each point is a mean over all individuals in a single replication population. Specialization is measured as the fraction of the population with a strong preference for environment A (90% chance or greater to reject environment B when encountered). K = 10,000, L = 100, μ_A_ = 5 × 10^−4^, μ_B_ = 5 × 10^−4^, μ_pref_ = 0.001, *p* = 0.25, and *c* = 0.25.

The examples in Fig. 2 illustrate how asymmetries in evolvability can lead to specialization, as well as the positive feedback that can lock in a narrow niche and prevent the emergence of a superior generalist genotype. These positive feedbacks are evident in the further skewing of the ratio of fitness after environmental preference evolves; as seen in Fig. 2B, fitness in environment B can decay below its initial level even as fitness in environment A is maximized. Note that such decay is expected because the deleterious rate is higher than the rate for beneficial mutations.

To further explore these results, I next focused on the quantitative effect of recombination. Figure 3 sweeps across a range of prevalences of environment B as well as intermediate levels of recombination. A population is classified as “specialist” if at least 90% of the individuals reject environment B with at least a 90% chance. Given that the model implements hard selection, frequency-dependent polymorphisms are not expected and this unidimensional measure is appropriate. However, these thresholds are obviously somewhat arbitrary, and increasing the required number of specialist individuals does have an effect on the outcomes, particularly at small values of *p* when selection on environment preference is weak (Supplemental Figure S1). Regardless, there is a clear overall pattern: recombination lowers the chance of niche reduction, allowing initial generalists to maintain that generalism as they adapt across a larger range of prevalences of the rare environment.

**Figure 3:**
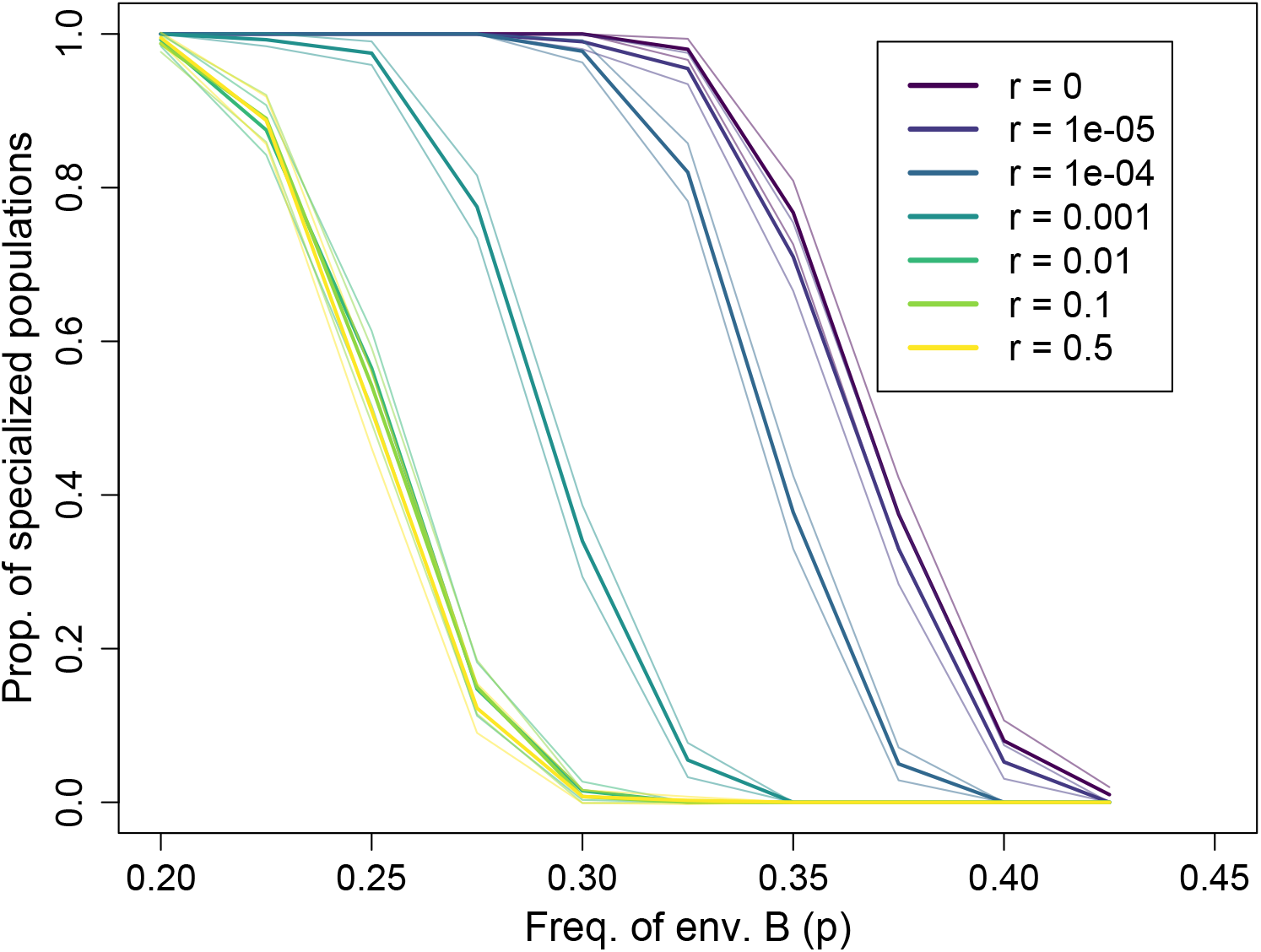
Evolved niche breadths across a range of values of p (*x-axis*) and r (*colors*). A population is classified as ‘specialists’ if at least 90% of individuals have at least a 90% rejection rate for the less common environment B. Thin lines indicate 95% confidence intervals based on the Normal approximation. Four hundred replicates were performed for each combination of parameters. As above, *K* = 10,000, *L* = 100, μ_A_ = 5 × 10^−4^, μ_B_ = 5 × 10^−4^, μ_pref_ = 0.001, and *c* = 0.25.

Figure 2 hints that specialization, once evolved, may persist after adaptation is over; I next validated this prediction for the set of simulations in Fig. 3. Among those populations (twenty-eight thousand populations across all treatments), over 99% could be classified as either specialist (containing at least 90% of individuals with a preference of at least 0.9 for environment A) or generalist (containing at most 10% of such specialist individuals) by generation 200,000. By the end of the simulations, at generation 400,000, every population could be classified as one or the other extreme. Moreover, none of the populations classified as generalist or specialist at the midpoint had changed their classification by the end. This stability indicates that while the process of adaptation was transient, its effects on preferences and therefore the utilized niche were long-lasting. These results also confirm that polymorphisms do not persist indefinitely under the conditions simulated here.

In Figure 3, recombination between the preference locus and the neighboring block of loci for performance in environment A also varies based on the stated *r* values. Additional simulations with free recombination between the preference locus and performance loci yield indistinguishable results (Supplemental Figure 2), confirming that the effect of recombination in Fig. 3 is caused by relieved interference between the genetic bases of performance in the two environments.

### The propensity for adaptive specialization is sensitive to determinants of evolvability

Schiffels et al.’s (2011) study of clonal interference predicted that the fixation rate of the largest-effect beneficial mutations sets the scale of emergent neutrality—greatly weakened effective selection on mutations with smaller effects. Much like the classical result that mutations with an *s* of much less than 1/*N* (in haploids where N_e_ equals *N*) are effectively neutral (Kimura 1968), Schiffels et al. update this relationship to predict that competing mutations in asexual populations are emergently neutral if *s* is much less than 1/*N* + *V*_drive_, where *V*_drive_ is the substitution rate of the large-effect, driver mutations. In assessing the causes and generality of asymmetric evolvability and the resulting evolution of specialism, I applied the results of Schiffels et al. (2011) to make three predictions for the model here. First, in asexual populations, increasing *N* or μ will worsen clonal interference and exacerbate the asymmetry between the rates of evolution in the common and rare environment. Second, the rate of fixation of beneficial mutations in the common environment will correlate with the asymmetry of adaptation and the propensity of specialism to evolve. Third, *V*_drive_ + 1/*N* will predict the degree of asymmetric evolvability by determining how much the rate of improvement in *w*_A_ causes improvement in *w*_B_ to lag.

Figure 4 shows that the product *N*μ does correlate positively in the absence of recombination with both asymmetry of adaptation rates (4A), and the probability for asexuals to evolve specialism (4C). Across experiments with different values of *N*μ, there is also the expected negative relationship between *V*_drive_ + 1/*N* and asymmetry (4B). However, increasing *N* and decreasing μ while maintaining their product, *N*μ, does result in greater asymmetry and probability of specialization, and a lower value of *V*_A_, contrary to the expectation derived from Schiffels et al. (2011). While *w*_B_/*w*_A_ is a good predictor of the probability to evolve specialism (4C), *V*_drive_ + 1/*N* does not capture the same information (4D). Note as well that the realized selection coefficient in generalists for adaptive mutations in environment B for asexuals is approximately the product of *p* and *s*, which is about 0.0025 for the results shown here. This is larger than any values of V_drive_ + 1/*N* achieved in these simulations; therefore, Fig. 4B&D should be seen as attempting to extrapolate Schiffels et al.’s prediction of an emergent neutrality threshold to predict evolutionary stalling for larger-effect mutations, and not as a test of their own claims for their result. Together, these results confirm the first two predictions derived from Schiffels et al. (2011), while illustrating that the broader phenomenon of evolutionary interference between traits is not yet fully understood.

**Figure 4:**
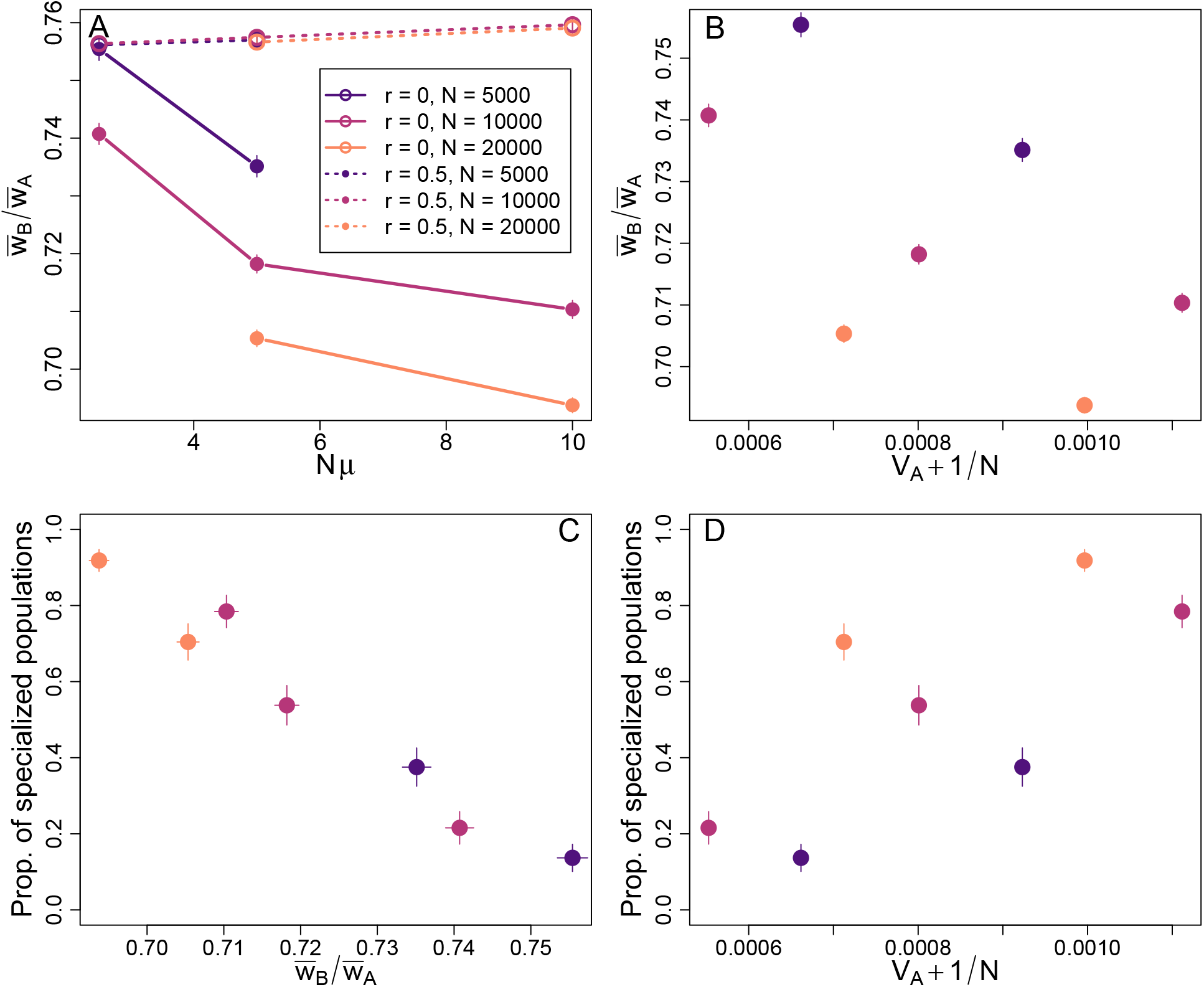
Asymmetry of adaptation, predicted neutrality threshold, and the fraction of specialists for combinations of *N* and μ. *Panel A*: With complete linkage, the ratio of fitness in environment B over environment A generally declines with increases in the product *N*μ (where μ = μ_A_ = μ_B_), indicating greater asymmetry of evolvability. Without linkage, there is no negative trend and sensitivity of the ratio to Nμ is very low. The ratio of fitnesses is the grand mean of the means of populations, each assayed when the mean of w_A_ crosses 0.8 (as determined by linear interpolation). *Panel B*: The ratio of fitnesses compared to the emergent neutrality predictor of substitution rate of adaptations in *w*_A_ plus the reciprocal of population size. Here and below, only the treatments with complete linkage are shown due to the lack of any substantial response of *w*_B_/*w*_A_ to *N*μ in populations without linkage. *Panels C and D*: Fraction of replicate populations that end their simulations as specialists, by the criteria applied above (e.g., Figure 3), plotted against the ratio of fitnesses and the emergent neutrality threshold. As above, *L* = 100 and *c* = 0.25. *w*_B_/*w*_A_ and *V*_drive_ are measured in simulations with μ_pref_ = 0 to avoid any confound caused by the evolution of specialism; fraction specialists is measured in separate simulations with μ_pref_ = 0.001. Values of *p* were chosen for *r* = 0 and *r* = 0.5 in which the fraction of populations evolving to specialists was approximately 0.5 when *K* = 10,000 and μ_A_ = 5 × 10^−4^, μ_B_ = 5 × 10^−4^; these values were *p* = 0.251 for *r* = 0 and *p* = 0.367 for *r* = 0.5, as estimated from the data in Figure 3.

Consideration of *V*_drive_ also motivates an exploration of how these results depend on the genetic architecture of fitness: specifically, how would *V*_drive_ and the probability of specialization change if each trait were determined by fewer loci of larger effect? Figure 5 shows that the probability of the evolution of specialism is generally robust to increasing *s* and decreasing *L* (such that *sL* remains equal to 1). Genetic architectures with fewer loci of larger effect adapt faster, as evident in Fig. 5B; this may be explained by the fact that the chance that a beneficial mutation escapes loss by drift increases approximately linearly for values of *s* in the range explored here. The evolution of specialism is well-predicted by the ratio of fitness gains and by *V*_drive_ + 1/*N* in populations with complete linkage, though not in populations with free recombination between the determinants of each trait. Populations with high values of *s* adapt much faster—when *r* = 0.5, populations achieve a mean *w*_A_ of 0.8 in about 56,000 generations when *s* = 0.01 but only about 2800 generations when *s* = 0.05, with a similar pattern of about 67,000 generations at s = 0.01 and 3300 at s = 0.05 for r = 0. The window of time for evolution of specialization is therefore likely to be much lower with large s, though why this shorter window might affect populations with recombination specifically (Fig. 5D) is not clear.

**Figure 5:**
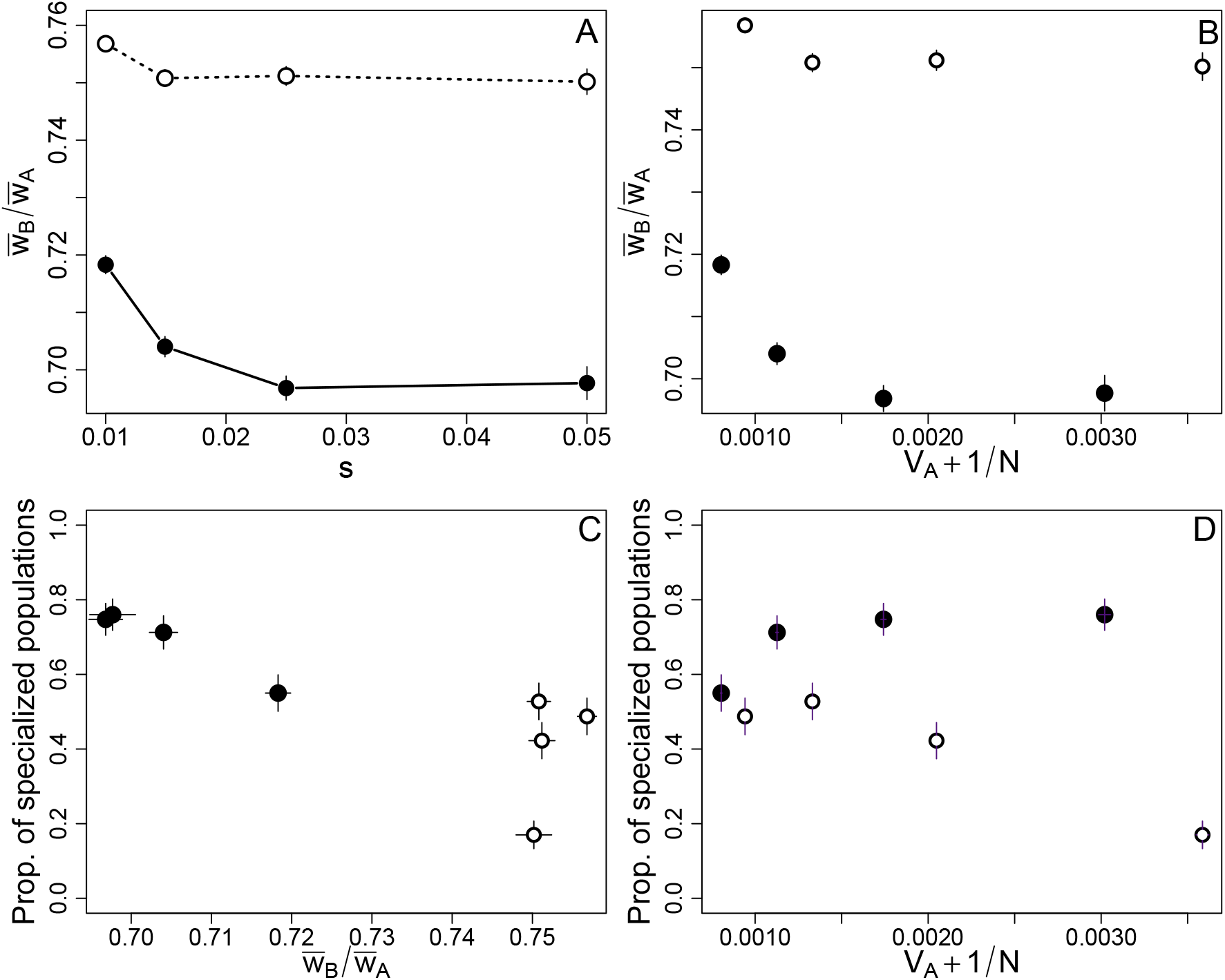
Asymmetry of adaptation, predicted neutrality threshold, and the fraction of specialists for combinations of *s* and *L. Filled circles*: *r* = 0; *open circles*: *r* = 0.5. For each tested value of *s, L* is reduced proportionally so that *sL* = 1; k, the number of beneficial alleles for initial genotypes, is also adjusted so that each treatment begins at approximately 50% of maximal fitness in each environment. *Panel A*: The ratio of fitness in environment B over environment A is maintained or slightly declines with increases in s (where *s* = s_A_ = s_B_). The ratio of fitnesses is the grand mean of the means of populations, each assayed when the mean of w_A_ crosses 0.8 (as determined by linear interpolation). *Panel B*: The ratio of fitnesses compared to the emergent neutrality predictor of substitution rate of adaptations in w_A_ plus the reciprocal of population size. *Panels C and D*: Fraction of replicate populations that specialized, by the criteria applied above (e.g., Figure 3), plotted against the ratio of fitnesses and the emergent neutrality threshold. *K* = 10,000, μ_A_ = 5 × 10^−4^, μ_B_ = 5 × 10^−4^, *c* = 0.25. *w*_B_/*w*_A_ and *V*_drive_ are measured in simulations with μ_pref_ = 0 to avoid any confound caused by the evolution of specialism; the proportion of specialized populations is measured in separate simulations with μ_pref_ = 0.001. Values of *p* were chosen for *r* = 0 and *r* = 0.5 in which the fraction of populations that specialized was approximately 0.5 when *K* = 10,000 and μ_A_ = 5 × 10^−4^, μ_B_ = 5 × 10^−4^; these values were *p* = 0.251 for *r* = 0 and *p* = 0.367 for *r* = 0.5, as estimated from the data in Figure 3.

### Negative density dependence restricts but does not eliminate adaptive specialization

In addition to search costs, organisms might also suffer negative effects of crowding that could inhibit specialization. As described in the *Model* section, the number of adults is regulated to a carrying capacity *K* before adults are choose their reproductive environment. Without habitat preferences, we expect (1-*p*)*K* adults to reproduce in environment A and *pK* in environment B. To implement a cost of high densities, I ran simulations in which fecundity in an environment was reduced if the actual number of adults in that environment exceeded the expectation without any preferences (see *Model--Density dependence*). For example, if, due to habitat preference, environment A contained 20% more adults than the neutral expectation of (1-*p*)*K*, then the fitness of all individuals in environment A was divided by a factor of 1.2.

This form of negative density-dependence should make specialization less profitable and reduce the likelihood of transitions to specialism. It does—the frequency of the rarer environment at which 50% of replication populations evolve into specialists (p_50_) is smaller with crowding effects (Figure 6), indicating that generalism is maintained for a broader range of conditions. This reduction in the chance of specialists to evolve appears approximately independent of recombination rate.

**Figure 6:**
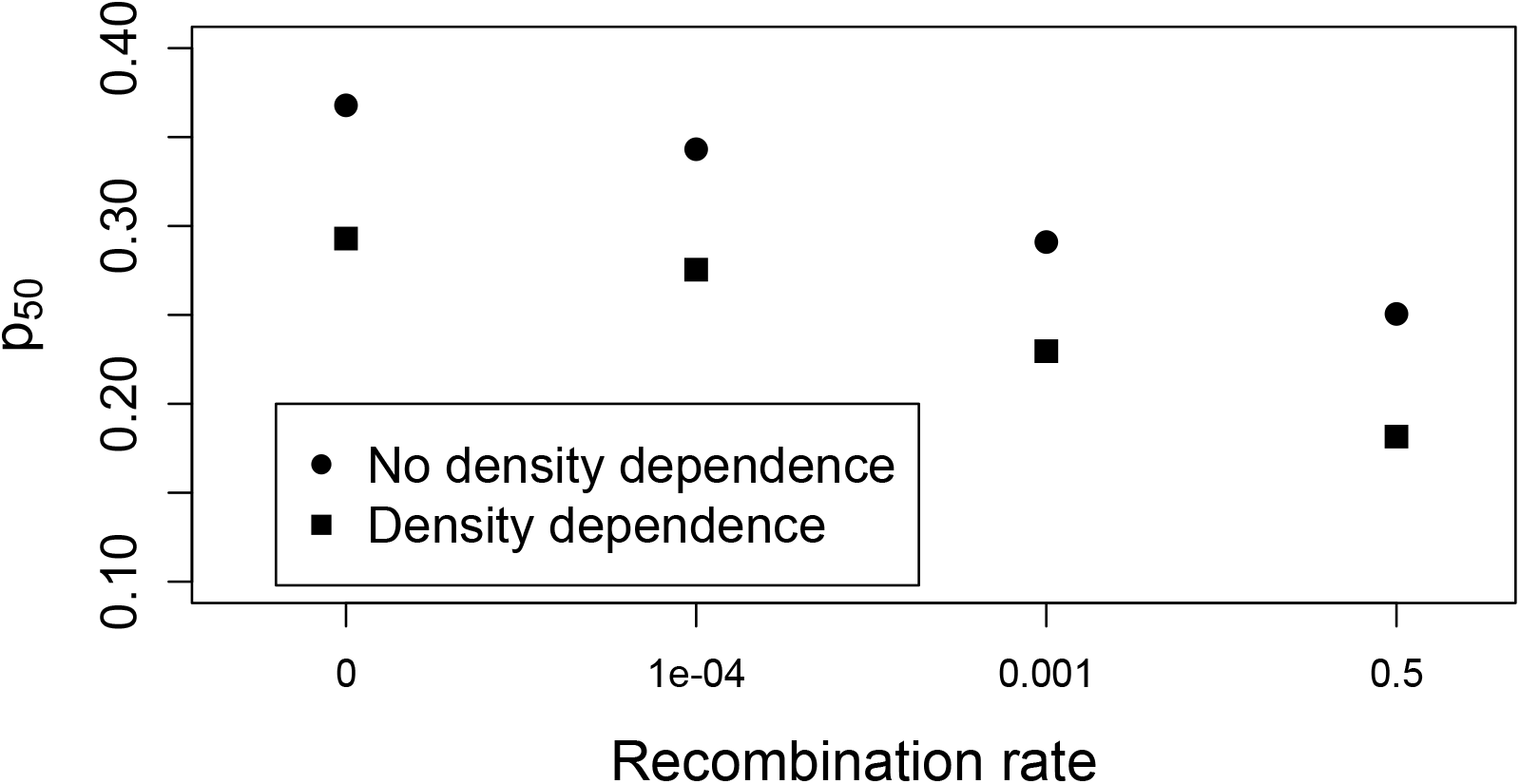
Value of p for which 50% of populations end the simulations as environment-A specialists (p_50_) with and without density dependence (squares vs. circles), across four values of the recombination rate. p_50_ was calculated by logistic regression; 95% confidence intervals were estimated by bootstrapping but omitted as they were well-approximated by the size of the plotting symbols. Data for the “no density dependence” treatment are the same as those summarized in Figure 3. As above, K = 10,000, L = 100, μ_A_ = 5 × 10^−4^, μ_B_ = 5 × 10^−4^, μ_pref_ = 0.001, and *c* = 0.25.

Negative density-dependent effects on fitness, as implemented here, could potentially lead to a frequency-dependent polymorphism of more- and less-specialized strategies. Such polymorphisms would render the focal summary statistic—fraction of populations where over 90% of individuals have over a 90% rejection rate of environment B—an incomplete picture of the evolving populations. However, after 500,000 generations of evolution, only two treatments out of the 48 combinations of *r* and *p* examined showed evidence of intermediate polymorphisms in preference—genotypes with prevalences over 10% and preferences between 0.1 and 0.9. With *r* = 0.5, such polymorphisms were found in five out of 244 replicates performed with *p* = 0.175 and nine out of 244 replicates with *p* = 0.2. While polymorphisms in preference may play a role during adaptive evolution, by the end of the simulations most populations were essentially monomorphic for preference—as was the case without density dependence.

## Discussion

The rejection of an unprofitable resource is a classical prediction (e.g., MacArthur & Pianka 1966), and evolved specialization in accordance with this prediction has been demonstrated empirically (Jasmin & Kassen 2007); this finding alone is not the key message of this paper. Instead, the focus here is on the embedding of this process of adaptive specialization within a context in which an environment becomes relatively unsuitable because performance in that environment fails to increase as quickly as performance in an alternative environment. My approach is more akin to models of habitat selection, in which populations may either adapt to better exploit unfavorable environments or to avoid them (Templeton & Rothman 1981; Castillo-Chavez et al. 1988; Rausher 1993; Feder & Forbes 2007; Ravigné et al. 2009). However, here the initial cause of a failure to adapt to an environment is that environment’s rarity, alongside the potential of ongoing adaptation to a more common environment to interfere with adaptation to the less common environment. Kawecki et al. (1997) found that mutation collapse of fitness in an unproductive environment depended only on that environment’s total contribution to the next generation. In contrast, the results here show adaptive collapse of fitness in environments with substantial initial, and potential, contributions to reproduction, but poor rates of improvement in that contribution.

Although rapid evolution is increasingly recognized as a contributor to ecological processes (e.g., Thompson 1998), much less is said about how the determinants of evolvability could explain patterns of biodiversity (but see Poisot et al. 2011). One argument is that regardless of whether adaptation is fast or slow, populations will still arrive at the same traits. This view is challenged by a primary role of rates of adaptations in a number of important scenarios: evolutionary rescue of populations fated for extinction (Bell 2017), rapid evolution in invasive species (Stapley et al. 2015), assembly of communities (Kremer & Klausmeier 2017), host-pathogen coevolution (Abrams & Matsuda 1997; Cortez & Weitz 2014; Hiltunen et al. 2014), and coexistence among competitors (Lankau 2011). The results here add another such scenario, in which a type of “race” between the performances of two adapting traits can decide which of two distinct, stable outcomes is reached. This work is therefore allied with a diverse set of models that examine which competition among distinct solutions to environmental heterogeneity (e.g., Bull 1987; Svardal et al. 2011; Tufto 2015). These models help to illustrate how subtle differences in rates of adaptation could matter, further motivating the idea that evolvability can act as an organizing framework for evolutionary biology (Wagner & Altenberg 1996; Pigliucci 2008).

The results here show one path to specialization without trade-offs, but do the data suggest that an alternative to trade-offs is required? Populations are often found to be adapted to their local conditions, but this fact alone doesn’t allow the inference that trade-offs are responsible for specialism. Kassen (2002) reviewed selection experiments in single environments and concluded that most showed evidence of negative genetic correlations for fitness across environments after selection for specialization. However, only about a third of these showed clear-cut evidence for general costs of adaptation—a decrease in fitness in unselected environments in parallel with specialization to the selected environment. Costs of adaptation may be caused by trade-offs, accumulation of mutations deleterious in unselected environments (Kawecki 1994), or a mix of both (e.g., Reboud & Bell 1997). These distinct causes can be teased apart by experiments that challenge some populations to evolve to multiple environments while others specialize on particular resources, or by examining parallel changes across replicate populations adapting to a single environment. Both approaches have yield mixed results. For example, experiments with VSV have repeatedly found that the virus adapts differently to distinct cell types, but can adapt to multiple cell lines simultaneously without apparent cost (Turner & Elena 2000; Remold et al. 2008; Smith-Tsurkan et al. 2010). A similar experiment that challenged *E. coli* to evolve to use glucose, lactose, or a combination found that generalists did suffer apparent trade-offs after an initial period of rapid improvement in both environments (Satterwhite & Cooper 2015). Lenski’s long-term evolution experiment with *E. coli* has been used to examine whether patterns in the decay of unselected functions indicates trade-offs or mere accumulation of unselected mutations—while an early analysis supported trade-offs (Cooper & Lenski 2000), a later re-examination supported mutational accumulation as the dominant factor (Leiby & Marx 2014). A yeast evolution experiment across eight environments showed a general pattern of specialization, but not necessarily trade-offs (Jerison et al. 2020); analysis of replicate populations showed a large role for stochastic forces in evolution. Chavhan et al. (2020) provide an empirical example in which costs of adaptation increase with population size, underlining the point that trade-offs are an outcome of evolutionary processes as well as a determinant of their course. Replicate experiments that yield trade-offs sometimes but not always are expected if possible mutations vary in their degree of antagonistic pleiotropy (Bono et al. 2017). Detailed experimental evolution approaches can quantify causation in costs of adaptation: for example, one study with digital organisms distinguished the effects of beneficial and neutral mutations on non-selected traits (Ostrowiski et al. 2007). These approaches for investigating trade-offs are more broadly applicable than just laboratory experiments; for example, repeated evolutionary loss of eyes and pigment in cavefish has been studied with the same dichotomy of trade-offs and mutation accumulation (Jeffrey 2009).

The rather uncertain relevance of trade-offs for specialization in evolution experiments echoes the debate over the causes of specialization in the field (Fry 1996). There are a number of reasons why negative correlations in performance across environments may not be patent even if trade-offs are important (Joshi & Thompson 1995). In some cases, evolution may have also ameliorated significant trade-offs by reducing overlaps in the genetic bases of conflicting traits (Rausher 1988). Futuyma & Moreno (1988) point out that negative performance correlations, even when found, cannot simply be interpreted as the cause of niche breadth because those same performance traits are products of evolution. This dilemma motivates the theoretical approach pursued here in which the values of the two environments, in terms of both initial and potential fitness, are equal; differences in performance and therefore the payoff of exploiting each environment can then only emerge by the interplay of the performance and preference traits.

The question of whether specialization can evolve without being driven by existing trade-offs has been more recently discussed in relation to microbes, particularly viruses (Remold 2012), but, earlier, was inspired by patterns seen in phytophagous insects (Fry 1996). In light of the results here, one puzzling characteristic of phytophagous insects is the existence of highly successful asexual lineages with very broad host ranges (reviewed in Gibson 2019). The work presented here does not model competition between sexual and asexual lineages, and therefore does not attempt to predict or explain this empirical pattern. However, there are several connections to be drawn between this model and relevant features of asexual, generalist insects that could guide future work. First, as summarized in Gibson (2019), asexual generalist insects typically have limited ability to disperse and choose habitats; this feature could be approximated in this model as a high value of *c*, the cost of searching. Second, generalists may suffer a reduced ability to evolve in response to change in any one aspect of their niche, but this evolvability deficit might be compensated for by their increased population size (Whitlock 1996). The results presented here rely on a sizeable gap between an organism’s current and optimal performance in each environment. Very large populations, whether sexual or asexual, may be able to keep pace with changing environments to such an extent that this performance gap does not arise. This reasoning parallels that expressed by Gibson (2019), who suggested that large populations with diverse hosts may benefit less from the effects of sex on evolvability, allowing more efficient asexual modes of reproduction to thrive. Experimental work by Hall et al. (2011) also highlights the idea that coevolutionary dynamics may induce different changes in niche breadth as compared to a single bout of adaptation to a static environment. Future work could explore how antagonistic coevolution with multiple hosts is shaped by the asymmetries in evolvability noted here.

The evolution of reduced niche breadth is often studied in models in which disruptive selection leads to partitioning of a niche by coexisting specialists (e.g., Roughgarden 1972), a process which is often linked with sympatric speciation (Doebeli & Dieckmann 2000). My emphasis here was on the evolutionary transition from generalism to specialism in a single lineage, not niche partitioning; hence, important topics for niche partitioning like frequency dependence were relegated to the background. Similarly, there is a rich history of modeling competition among generalists and specialists, which shares overlapping concerns but foregrounds ecological factors like temporal heterogeneity, frequency-dependent selection, and adaptive behavior that allow coexistence (Wilson & Yoshimura 1994). Future work could build on the insights illustrated here to couple evolvability with a more fully realized ecological model. Similarly, the absence of trade-offs is not a requirement for the results described here; trade-offs are excluded to isolate focus on the role of evolvability. Future work could readily add trade-offs to the framework explored here to understand how genetic limitations on generalist fitness interact with evolvability differences between traits. The model here also does not consider philopatry or other scenarios in which a lineage experiences one environment for many sequential generations; this choice distinguishes the focus from models of local adaptation and specialization like Ronce & Kirkpatrick (2001). Future work could look at asymmetric evolvability across traits in ecological scenarios with more limited migration between environments.

This model relies on positive feedbacks between habitat preference and performance, mediated by the effect of each on the intensity of selection for the other. Joint evolution of this pair of traits has a rich history, as summarized above, but other traits can show the same type of positive interdependence—for example, feeding efficiency in a forager changes the consumption of a given resource, and improvements in feeding efficiency and conversion efficiency of the same resource therefore interact synergistically (Vasconcelous & Rueffler 2020). More broadly, synergistic epistasis among aspects of fitness might lead to accelerated adaptation to one part of a niche, setting the stage for specialism to evolve. Based on this speculation, future work might profitably extend the model explored here to include more explicit links between traits, ecology, and fitness.

These results leverage the findings of Schiffels et al. (2011), and later work by Gomez et al. (2020) and Venkataram et al. (2019), to illustrate that asexual adaptation is not just slower than adaptation with recombination, but differs in other significant aspects. Other models have predicted differences in the nature of adaptation in asexual versus sexual organisms, such as difference in epistasis among fixed mutations (Livnat et al. 2008) and, as discussed above, asexuality is linked with extreme generalism in some insects (Gibson 2019). Still, there has not been a broad, synthetic effort to understand how sex and other important determinants of evolvability shape not just the rate of evolution, but bias adaptation toward distinct phenotypes, niches, or life-histories. It is hoped that this work will help push toward such an effort, and start to fully realize the goal of integrating evolvability with our understanding of the forces shaping the niche.

## Acknowledgements

[omitted during double-blind review]

---

**Supplemental Figure 1:**
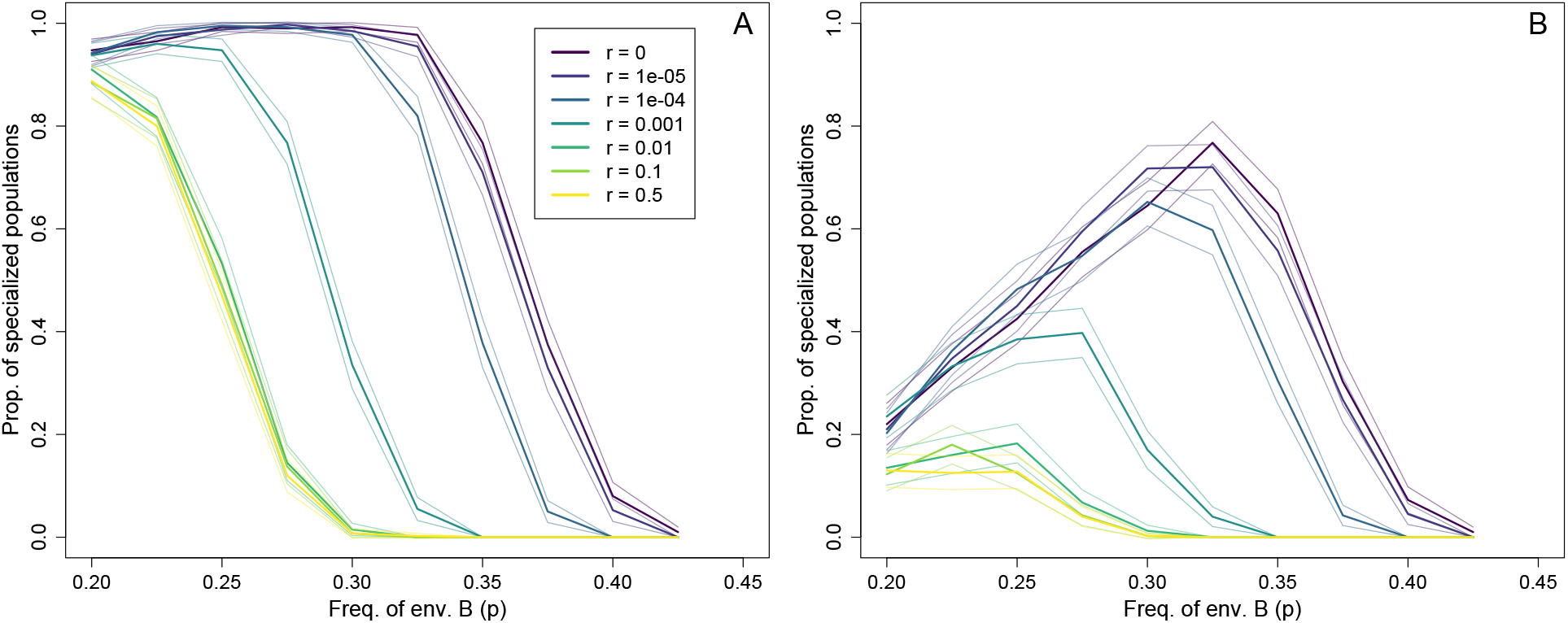
Evolved niche breadths across a range of values of p (*x-axis*) and r (*colors*). A population is classified as ‘specialists’ if at least 95% (*panel A*) or 97.5% (*panel B*) of individuals have at least a 90% rejection rate for the less common environment B. Thin lines indicate 95% confidence intervals based on the normal approximation. Four hundred replicates were performed for each combination of parameters. As above, *K* = 10,000, *L* = 100, μ_A_ = 5 × 10^−4^, μ_B_ = 5 × 10^−4^, μ_pref_ = 0.001, and *c* = 0.25.

**Supplemental Figure 2:**
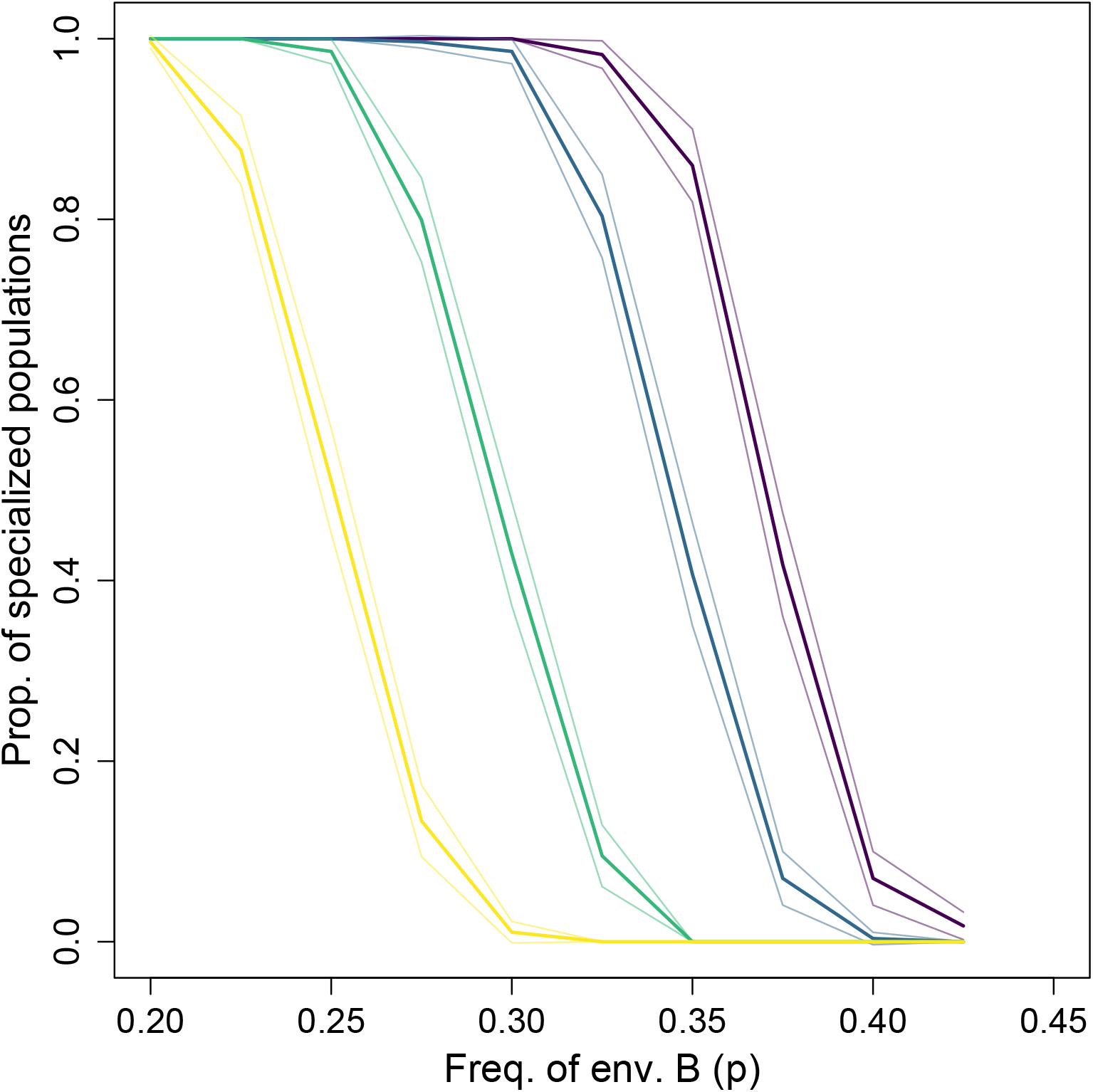
Evolved niche breadths across a range of values of p (*x-axis*) and r (*colors*). In contrast to Fig. 3, here *r* = 0.5 between the preference locus and the set of loci for performance in A. A population is classified as ‘specialists’ if at least 90% of individuals have at least a 90% rejection rate for the less common environment B. Thin lines indicate 95% confidence intervals based on the normal approximation. Four hundred replicates were performed for each combination of parameters. As above, *K* = 10,000, *L* = 100, μ_A_ = 5 × 10^−4^, μ_B_ = 5 × 10^−4^, μ_pref_ = 0.001, and *c* = 0.25.

